# Mistranslation protects against lifespan reduction due to mating in *Drosophila melanogaster* females

**DOI:** 10.64898/2025.12.01.691645

**Authors:** Luana Branco, William Yeung, Joshua R. Isaacson, Amanda J. Moehring

## Abstract

Accurate translation of genes into proteins is critical to organism fitness, and errors in this process are usually detrimental and cause proteotoxic stress. Mistranslation occurs when the amino acid that is incorporated into the nascent polypeptide chain does not match what is dictated by the genetic code. Valine-to-serine (V→S) and threonine-to-serine (T→S) mistranslating models of the fruit fly *Drosophila melanogaster* have demonstrated a surprising, sex-specific increase in virgin female longevity. We predict that the added stressor of reproduction would eliminate this mistranslation-induced lifespan increase since females prioritize reproductive tissues over somatic tissues, and proteotoxic stress would therefore lead to higher protein damage in the soma of mated females. We measured the impact of reproduction on V→S and T→S mistranslating *D. melanogaster* by measuring longevity, egg laying, and fecundity. Counter to our prediction, both V→S and T→S mistranslation led to a sex-specific increase in mated female longevity compared to non-mistranslating controls. Additionally, the risk of death decreased for mated females with mistranslation, beyond the pure additive benefits of mistranslation alone. These effects could not be explained by reduced egg laying or fertilization rates in mistranslating females. Thus, we find that the added proteotoxic stress caused by mistranslation does not exacerbate the detrimental effects of reproduction and instead can ameliorate lifespan decreases due to female reproduction.

## Introduction

Reproduction has high physiological costs and investing in reproduction can shorten lifespan (reviewed in Maklakov and Immler 2016). According to the Disposable Soma Theory, this lifespan reduction is due to the costly allocation of energy towards gamete production at the expense of somatic maintenance (Kirkwood 1977a, Kirkwoord 1997b; Kirkwood and Holliday 1997; reviewed in Flatt 2011). In *Drosophila melanogaster*, oogenesis and embryogenesis are tightly regulated processes with high energetic and nutritional needs. High metabolic investment is required from females to produce oocytes containing the yolk proteins, lipids, and carbohydrates necessary to undergo future embryogenesis (Gutzeit *et al*. 1994; Buszczak *et al*. 2002; Sieber *et al*. 2016). To support high metabolic needs, a high protein diet is often favoured by mated females compared to males and unmated females (Zheng *et al*. 2025). In addition to high protein intake, female *D. melanogaster* invest energy into maintaining proteostasis in germline tissue at the expense of somatic tissue, prioritizing offspring investment over parental longevity (Fredriksson *et al*. 2012). The prioritization of germline protein quality control over somatic tissue in mated females should therefore lead to a more pronounced impact of proteotoxic stress on somatic traits, such as lifespan, compared to the same traits in males and unmated females.

Inaccurate translation of mRNA into proteins can produce misfolded proteins and disrupt proteostasis, compromising organismal health and fitness (Santos *et al*. 2018; Tavares *et al*. 2021). Mistranslation, defined as the erroneous incorporation of non-cognate amino acids into a nascent polypeptide chain, has been primarily studied *in vitro* using single-cell models (*Escherichia coli, Saccharomyces cerevisiae*, human cell lines: e.g., Berg *et al*. 2021; Berg *et al*. 2022; Davey-Young *et al*. 2024; Samhita *et al*. 2020, Zhang and Ling 2025). In these studies, mistranslation causes defects in protein production, increases in protein aggregation, proteotoxic stress, and growth defects (Paredes *et al*. 2012; McDonald *et al*. 2025; Tennakoon *et al*. 2025). Mistranslation is usually detrimental but can provide benefits in certain circumstances. In yeast and bacterial species, alteration of the genetic code can confer an evolutionary advantage in evading host immune responses and developing antibiotic resistance (Pezo *et al*. 2004; Bacher *et al*. 2007; Silva *et al*. 2007; Vargas-Rodriguez *et al*. 2021; Schuntermann *et al*. 2023). Additionally, mistranslation has been shown to protect cells against oxidative stress, increase cell survival under UV-induced DNA damage, and upregulate the yeast longevity gene *PNC1* (Netzer *et al*. 2009; Silva *et al*. 2009; reviewed in de Pouplana *et al*. 2014; Fan *et al*. 2015; Samhita *et al*. 2020). Despite mistranslating tRNA variants being common in humans (Lant *et al*. 2018; Berg *et al*. 2019; Hasan *et al*. 2023; Tennakoon *et al*. 2025) and mistranslation being implicated in disease (Bloom *et al*. 2006; Kalapis *et al*. 2015; Lant *et al*. 2018; Santos *et al*. 2018), our understanding of how it affects various tissues in multicellular organisms is limited by a scarcity of *in vivo* research using multicellular models.

Recent studies in *D. melanogaster* revealed that variant transfer RNA (tRNA) genes that induce valine-to-serine (V→S) or threonine-to-serine (T→S) mistranslation increased longevity in unmated female flies (Isaacson *et al*. 2024b). The underlying mechanism for this increase in longevity is unknown but may be due to mistranslation activating stress response pathways, such as those affecting the cell cycle and DNA repair (Isaacson *et al*. 2024a), causing a general protective effect against other stressors that could impact longevity. However, reproduction would shift cell maintenance towards reproductive tissues at the expense of somatic tissues, and gamete production and maturation require increased protein synthesis activity and reallocation. We therefore predict that mistranslation’s beneficial effect on unmated female lifespan will be removed when females are reproducing, and that their lifespan will be shorter than controls due to the combined detrimental effects of prioritizing germline tissue maintenance at the expense of somatic tissue, proteotoxic stress from mistranslation, and energetic costs of reproduction (Salmon *et al*. 2001; Harvanek *et al*. 2017).

To test this prediction, we assess the effects of mating and mistranslation on *D. melanogaster* lifespan. We found that reproduction decreases longevity in females, as expected, but that tRNA-induced mistranslation buffers its negative impact, counter to prediction. Mistranslation does not affect egg laying or fertilization, and therefore the increase in longevity in mistranslating mated females is not due to failure of basic reproductive output. Hence, mistranslation increases lifespan even in the presence of additional proteotoxic burdens placed on females because of mating.

## Methods

### Fly husbandry and stocks

All flies were housed at 24°C, 75% relative humidity (RH), 14:10 light:dark (L:D) (longevity assay) and 25°C, 80% RH, 12:12 L:D (egg laying and fecundity assays). Flies were housed on Bloomington standard cornmeal agar food media recipe (Bloomington Drosophila Stock Center). Mistranslating flies were created using a tRNA model first developed in *S. cerevisiae* (Berg *et al*. 2017; Berg *et al*. 2019). In *D. melanogaster*, a tRNA^Ser^ containing the anticodons for valine (AAC) or threonine (AGU), and a G26A secondary mutation, were integrated into a set landing site on the second chromosome (Isaacson *et al*. 2024b). These tRNAs will recognize valine or threonine codons whilst remaining serylated, resulting in valine-to-serine (V→S) or threonine-to-serine (T→S) mistranslation. Mistranslation was confirmed via mass spectrometry (Isaacson *et al*. 2024b). Mistranslating stocks had FRT sites flanking the mistranslating tRNA sequence, allowing for the removal of the tRNA gene in the presence of flippase to create control flies (Gronostajski and Sadowski 1985; Isaacson *et al*. 2024b). The control flies have an identical genetic background to the mistranslating flies, with the only difference being the removal of the mistranslating tRNA. The genotype of the flies with the mistranslating tRNA is: *w^1118^; P{CaryP}-attP40^wmw+=pattB-tRNA^/CyO; MKRS/TM6B*. The genotype of the control flies is: *w1118; P{CaryP}-attP40^wmw+=pattB-FLP-out^/CyO; MKRS/TM6B*. Wildtype *Canton S* males were used as a control mate for the egg laying and fecundity assays.

### Longevity assaysh

Longevity assays were conducted as described in Isaacson *et al*. (2024b). We collected equal numbers of virgin males and females of each mistranslating line (V→S and T→S) and their controls under light CO_2_ anaesthesia within 8 hours of eclosion, removed any with visible signs of deformity (following the protocol in Isaacson *et al*. 2022), separated by sex, and placed in new food vials. For each mistranslating line, virgin flies were divided into replicates of eight vials each, with each vial within the replicate containing 4-8 (V→S) or 4-6 (T→S) flies (Fig. 1; Supplemental figure S1). Four vials contained unmated flies, containing a single sex (male or female) and genotype (mistranslating or control). Four vials contained mated flies, containing equal numbers of males and females, with the total number being equal to that for the unmated flies, for each genotype (mistranslating or control). All vials set up on a given day contained the same number of flies per vial across all groups to minimize environmental effects (Austin *et al*. 2014). For example, if on a given day there were 4 unmated flies per vial, then there would be two corresponding vials that each contain 2 mated males and 2 females (Supplemental figure S1); if on another day there were 6 unmated flies per vial, then there would be two mated vials that each contain 3 males and 3 females. Flies that escaped or were killed during transfer were censored from the observation. In total, 82 flies from each group were tested from the V→S cohort and 80 were tested from the T→S cohort (the breakdown of vials and flies is listed in Supplemental Table S1). Every 3 days, flies were transferred to new food and deaths were recorded.

**Figure 1.**
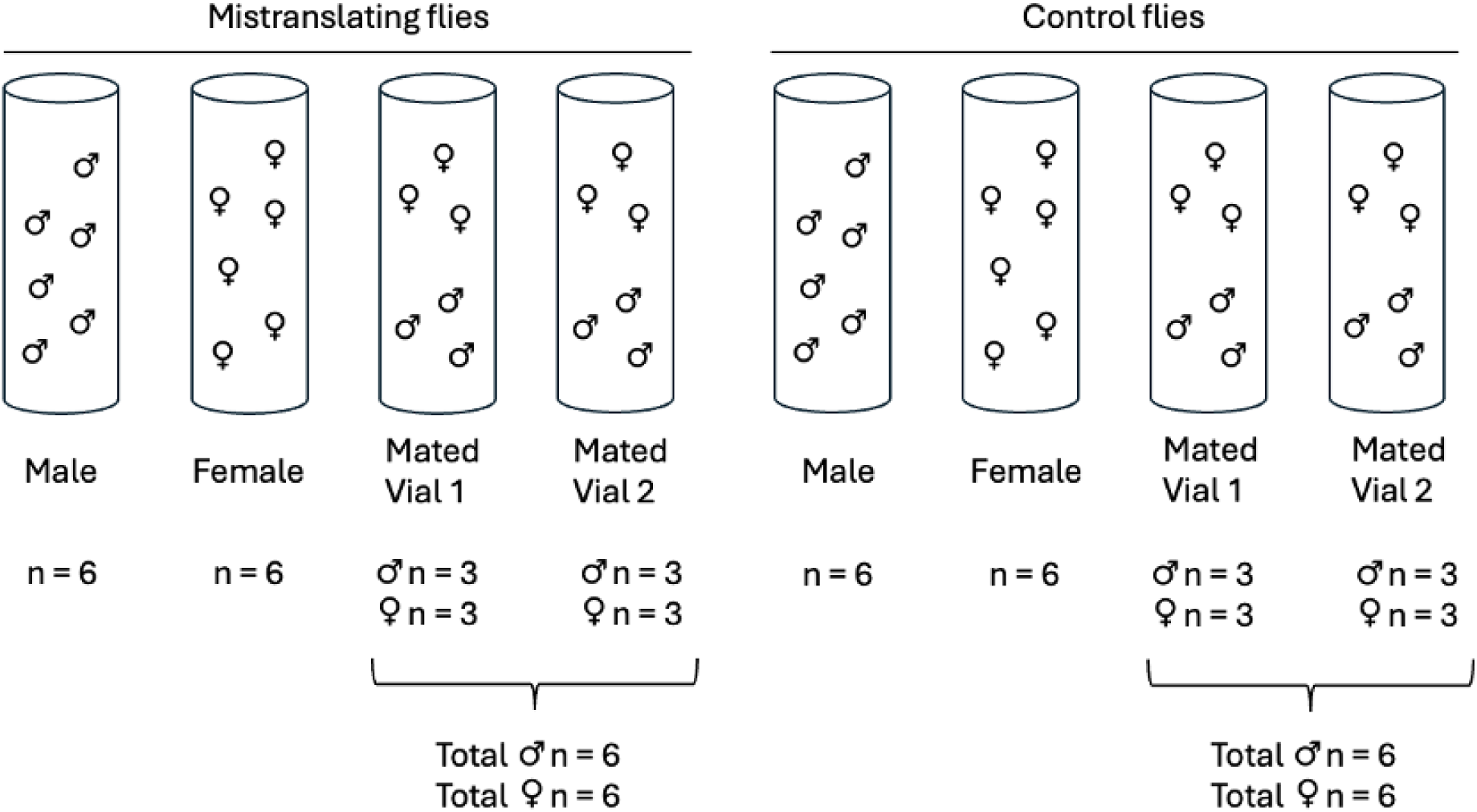
Longevity assay setup on a given day. Each male is depicted in the vials by the “♂” symbol and females are depicted by the “♀” symbol. The number of each fly in each vial is displayed beneath the vial. For a given set, vials contained a total of 4, 6, or 8 flies, with all vials within that set having the same number. The example of a total of 6 flies is shown; for examples of 4 and 8 flies see Supplemental Figure S1.

### Egg laying and fertilization assays

Virgin wildtype *Canton S* (*CS*) males and virgin mistranslating and control females were collected under light CO_2_ anaesthesia and separated by sex. Flies were aged 3-7 days prior to being placed in single mating pairs and observed for up to 2 hours until mating occurred; unmated pairs were discarded. We transferred females 20-30 minutes post-mating to a fresh 35 x 10 mm grape juice plate (Nouhaud *et al*. 2018) containing 10-30 mg of 1:1 ddH_2_O and yeast mixture added to the surface. We allowed females to lay eggs for approximately 24 hours, transferred females to a new plate, and counted the number of eggs each female laid up to 3 hours past lights on (Nouhaud *et al*. 2018). To assess eggs that were fertilized from sperm coming from short-term sperm storage versus long-term sperm storage, we assessed egg laying on days 1-4 and 7-10 post-mating, respectively (Dhillon *et al*. 2020). Females were transferred to new grape juice plates every 24 hours for the days that egg counting was performed; in between the short-term and long-term sperm storage observation, female flies were kept on Bloomington standard cornmeal agar food media. For V→S genotypes, we assessed egg laying for 8 control females and 10 mistranslating females; for T→S genotypes, we assessed 12 control females and 11 mistranslating females. To confirm fertilization of eggs in both short-term and long-term sperm storage periods, we examined the fertilization status of the eggs laid by mistranslating and control flies. We considered eggs to be fertilized if they showed visible cellularization when assessed under a stereo microscope (Lecuit and Wieschaus 2000). For V→S genotypes, we assessed 20 eggs chosen at random across all the females for their fertilization status. To randomize the egg selection, we numbered all eggs and used a random number generator to determine which 20 would be selected. For T→S females, we assessed all eggs from all females for their fertilization status.

### Statistical analyses

We performed the statistical analysis and graphing in RStudio version 2024.12.0+467 running R 4.3.2 unless otherwise noted. We created longevity Kaplan-Meier curves (*survival* and *survminer* packages in R) as described in Isaacson *et al*. (2024b) and adjusted using the Benjamini-Hochberg (BH) false discovery rate (FDR) procedure (Supplemental Table S2, Benjamini and Hochberg 1995). To analyze the interaction effects between mistranslation and mating, we used Cox proportional hazard models (*survival* package in R). We included initial vial population size as a covariate to control for crowding effects. Additionally, we tested for random effects across individual vials by including vial ID as a frailty term; however, vial ID had negligible effects in both the V→S and T→S models (frailty variance estimate = 5x10^-7^ and 5x10^-3^, LRT *P* = 0.88 and 0.27, respectively) and so was excluded from downstream analyses. When verifying proportional hazards assumptions, we found that the Schoenfeld residual plot of mistranslation exhibited time-dependency; thus, we included a time-transformation term for mistranslation as an additional covariate. We adjusted *P*-values for multiple testing of covariates across Cox models using the BH FDR procedure, as above for longevity. Lastly, we calculated the confidence intervals and standard deviations of survival medians in Microsoft Excel Version 16.101.1.

Egg laying analysis was performed using linear mixed-effects (n*lme* package in R) to assess the effects of mistranslation, sperm storage period, and their interaction on egg laying behaviour. We used a square-root transformation to normalize the egg laying data and also included fly ID as a random intercept to control for random effects from within-fly variability. We then obtained the estimated marginal means and their standard errors (*emmeans* package in R) for visualization. We used a Wilcoxon test to analyze long-term and short-term egg laying when either pooled or separated by sperm storage periods. All data figures were generated in R using *ggplot2*.

## Results

We assayed *D. melanogaster* that express either a tRNA that mistranslates valine-to-serine (V→S) or threonine-to-serine (T→S; Isaacson *et al*. 2024b) to determine the effects of mistranslation on lifespan when flies are unmated *vs* mated (Figure 1, Supplemental Figure S1, Supplemental Table S1). We first compared the longevity of mated and unmated flies to confirm the known effect of mating on reducing female lifespan (Kirkwood and Holliday 1997; Fredriksson *et al*. 2012; Koliada *et al*. 2020). As expected, female flies containing a V→S or T→S mistranslating tRNA and their associated controls had significantly decreased longevity due to mating (*P*<0.0001 for all comparisons; Table 1, Figure 2, Supplemental Table S2). In contrast, males with a V→S or T→S mistranslating tRNA and their associated controls did not have a significant decrease in lifespan due to mating (Table 1, Figure 2, Supplemental Table S2). This aligns with the observation of Koliada *et al*. (2020) that reproduction causes a stronger detrimental effect in female compared to male lifespan.

**Figure 2.**
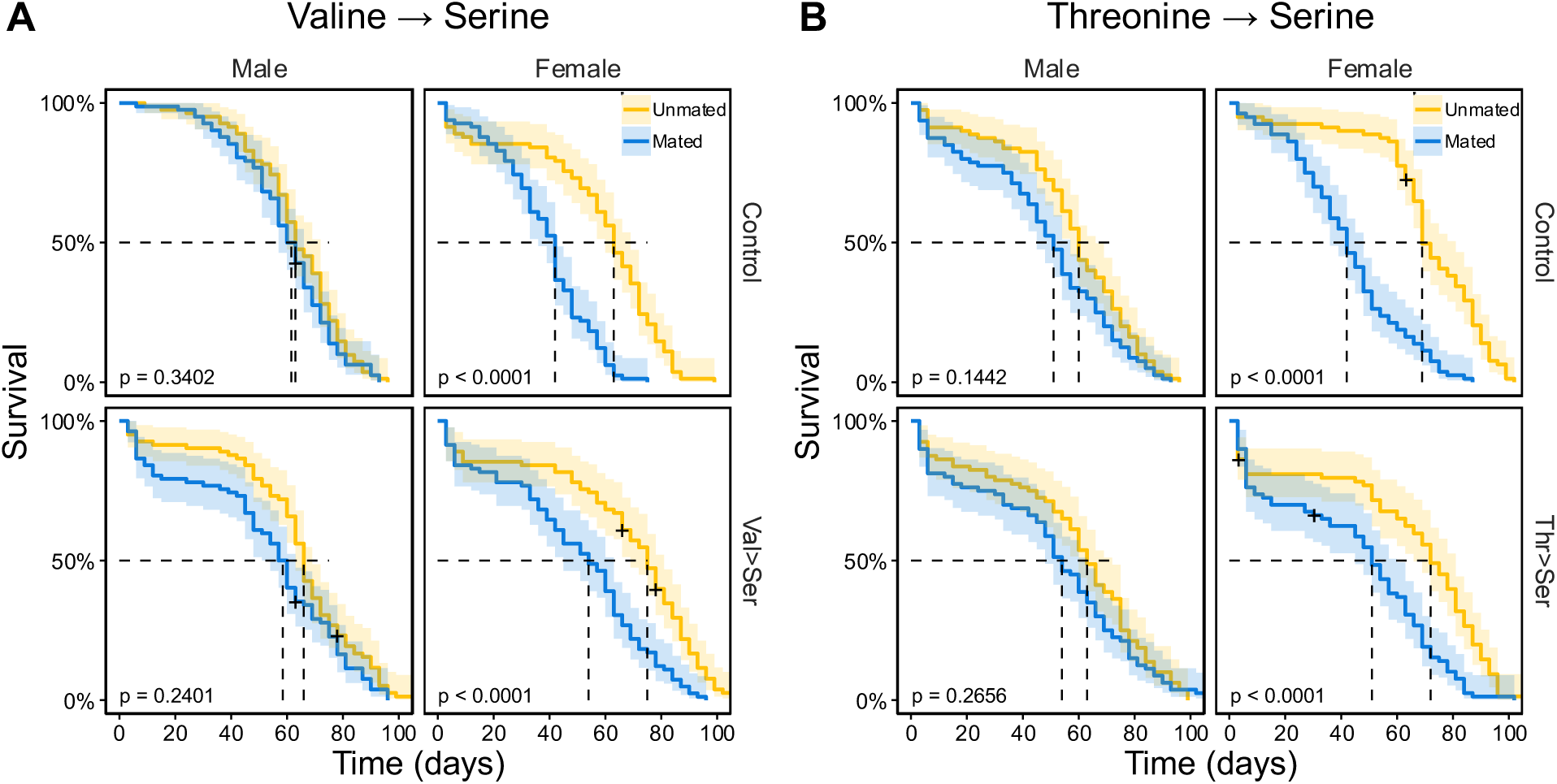
Survival of unmated (blue) *vs* mated (yellow) male or female flies for: a) V→S and b) T→S mistranslating males and females and corresponding controls. Shaded regions represent the 95% confidence intervals and plus (+) symbols represent flies that were censored from the observation (i.e., loss during transfer). 50% survival is displayed by dashed lines.

**Table 1.**
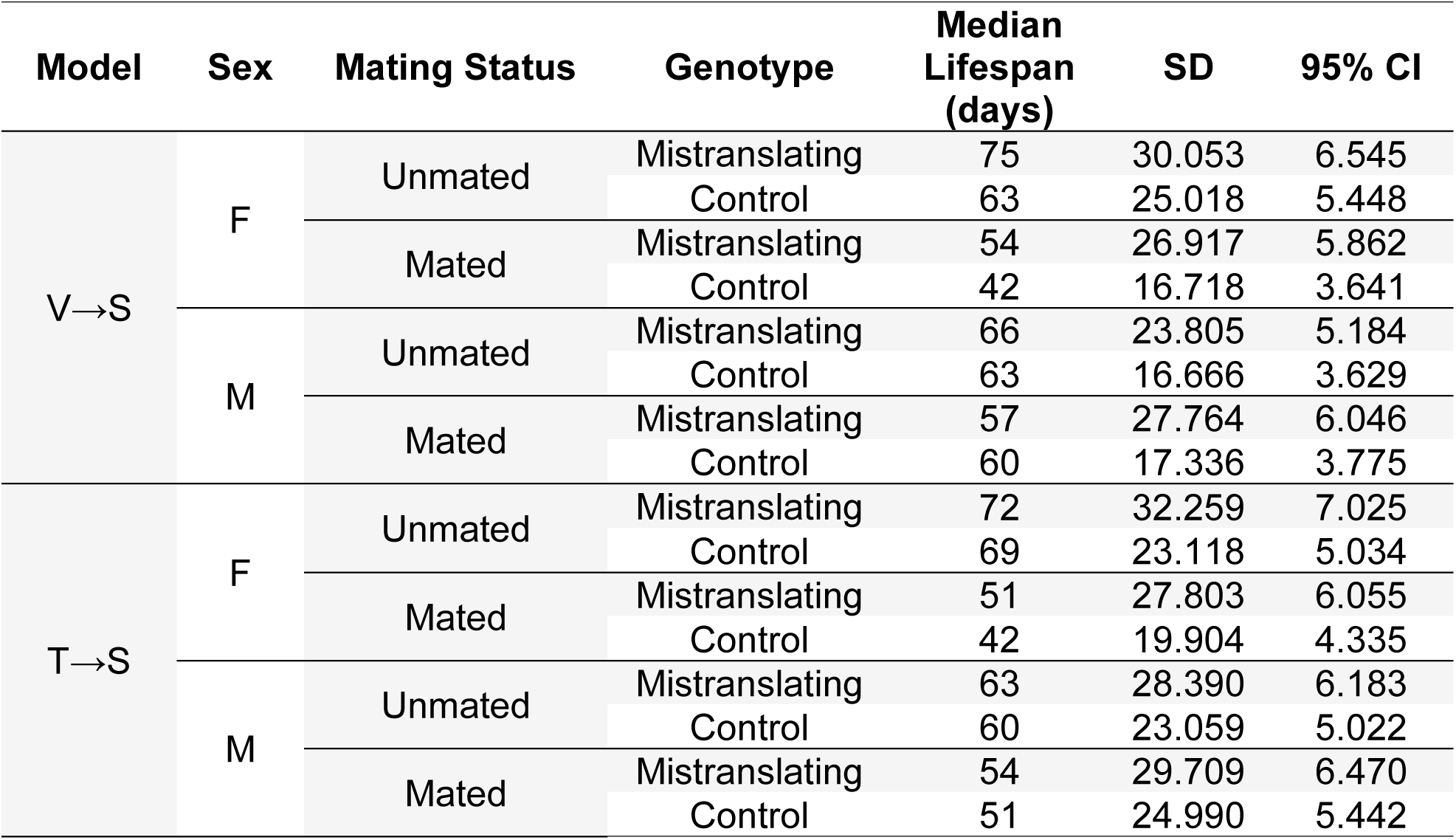
Median lifespan for each genotype tested according to mistranslation model, sex, and mating status.

We next assessed whether the lifespan improvement previously seen for mistranslating unmated females (Isaacson *et al*. 2024b) is eliminated when females divert investment into reproduction, if any observed effects are also present in males, and if effects are present for both types of mistranslation. To this end, we compared the survivorship of mistranslating lines and their controls for each sex for V→S and T→S mistranslation (Table 1, Figure 3, Supplemental Table S2). V→S mistranslation increased the longevity of both unmated (*P*<0.0001) and mated (*P*<0.0001) females compared to their controls (Figure 3a). These results indicate that mistranslation from valine to serine leads to an increase in the longevity of female *D. melanogaster*, regardless of mating status. In contrast, mistranslation from V→S did not have a significant impact on unmated and mated male longevity (Figure 3a), further demonstrating that V→S mistranslation has sex-specific effects. Unlike what was seen in V→S, mistranslation from T→S in unmated females and males did not have a significant effect on longevity when compared to their respective controls (Figure 3b). However, similar to what was seen for V→S, mistranslation from T→S in mated females resulted in a significant increase in longevity when compared to control females (*P*=0.0011), while males had no change in lifespan (Figure 3b). This indicates that T→S mistranslation, like V→S, causes sex-specific increased longevity in mated females.

**Figure 3.**
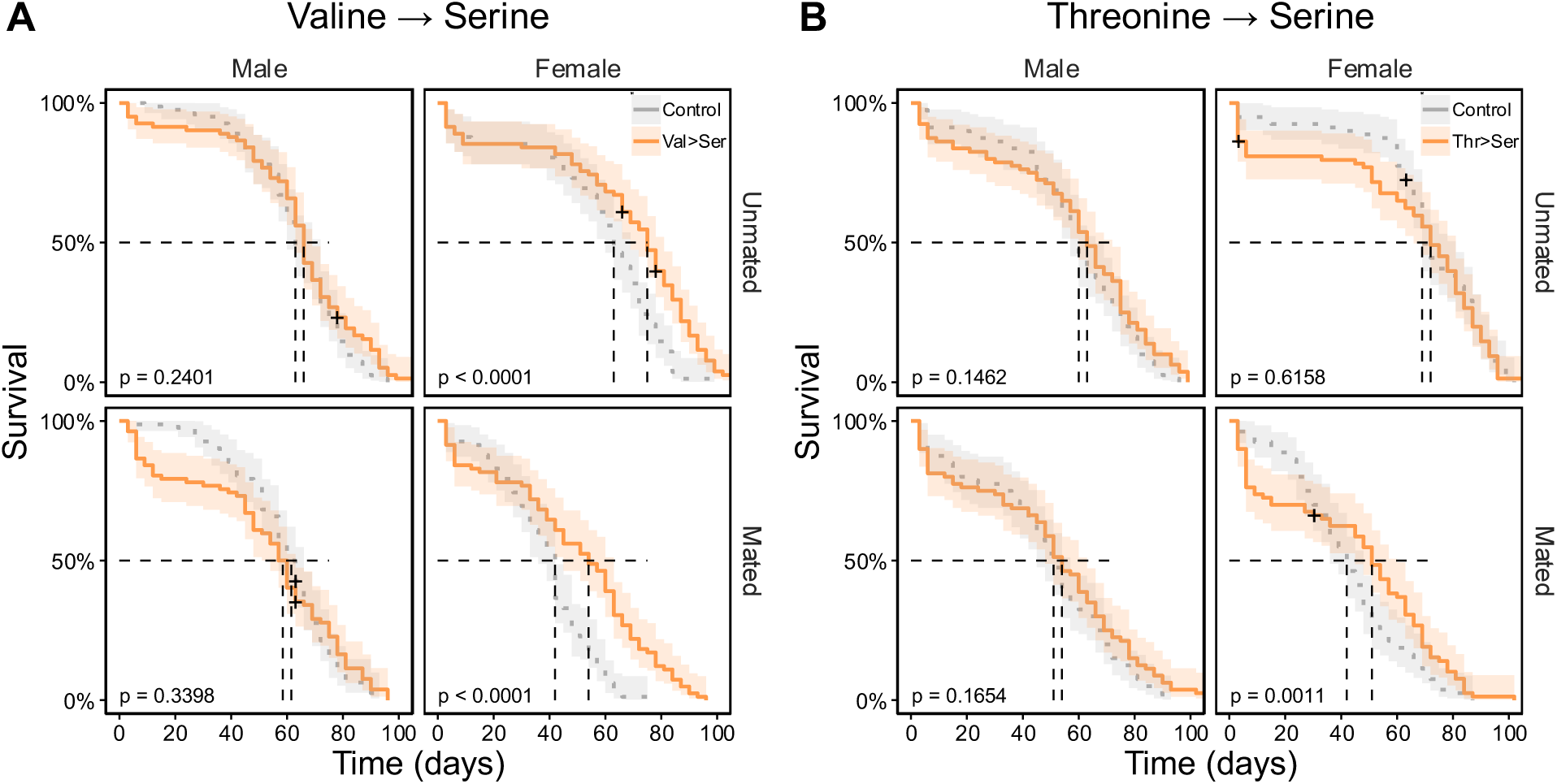
Survival of mistranslating flies (orange) *vs* their respective controls (gray) for male or female a) V→S or b) T→S mistranslation while mated or unmated. Shaded regions represent the 95% confidence intervals and plus (+) symbols represent flies that were censored from the observation (i.e., loss during transfer). 50% survival is displayed by dashed black lines.

We next wanted to ask if the presence of mistranslation ‘rescues’ mated female longevity to be the same length as for mated males; in other words, if it is sufficient to remove the sex-specific impact of mating on longevity. To determine the baseline effect without mistranslation in our flies, we compared the longevity of controls. Unmated control females and males did not differ in lifespan, while mated V→S and T→S control female lifespan was significantly shorter than mated control male lifespan (*P*<0.001, *P*=0.0049, respectively; Table 1, Figure 4, Supplemental Table S2). In other words, mating reduces lifespan in control females compared to males, as expected. When females are unmated, mistranslation significantly increases female lifespan to be greater than males in the presence of both V→S (*P*=0.0079; Figure 4a) and T→S (*P*= 0.014; Figure 4b) mistranslation. When females are mated, mistranslation does not increase lifespan to be greater than males, but does ‘rescue’ it to be the same as males, as we did not see a significant difference between mated female and male lifespan for either V→S (Figure 4a) or T→S (Figure 4b)

**Figure 4.**
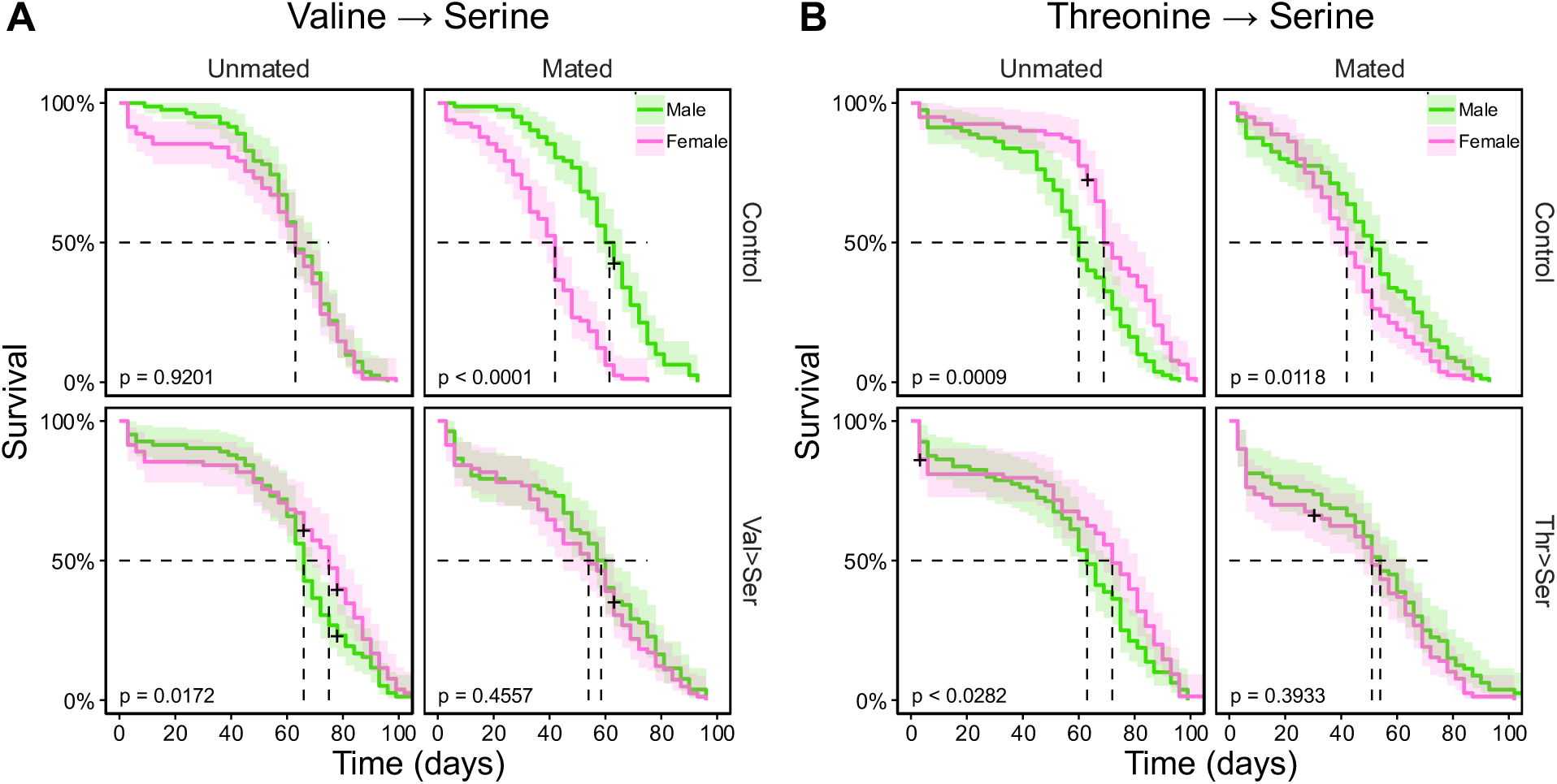
Survival of females (pink) *vs* males (green) for a) V→S and b) T→S mistranslating flies and controls while mated or unmated. Shaded regions represent the 95% confidence intervals and plus (+) symbols represent flies that were censored from the observation (i.e., loss during transfer). 50% survival is displayed by dashed lines.

mistranslation. Therefore, while mistranslation did not extend the lifespan of mated females beyond that of mated males, mistranslation prevented mated female lifespan from being significantly shorter than mated male lifespan, as is seen with their control counterparts. In other words, mistranslation buffers against the typical lifespan reduction seen in mated females (Burger and Promislow 2004). These results indicate that the presence of mistranslation provides an advantage to female longevity, regardless of mating status.

It is possible that the ‘buffering’ effect is simply an additive increase in longevity that is present regardless of mating status. To assess if the effects of mating and mistranslation on longevity are additive *vs.* synergistic, we analyzed predicted risk (of death) using Cox proportional hazard models. In alignment with the outcomes of our survival curve analyses, we found that mating increased risk in both V→S females (*P*<0.0032; Supplemental Table S4) and T→S females (*P*<0.0001; Supplemental Table S5). Our analysis also showed that the time-independent mistranslation covariate increases risk in both V→S females and T→S females (*P*<0.0001 for both), but that the time-*dependent* mistranslation covariate decreases risk in both (*P*<0.0001 for both). We calculated the timepoint at which the time-dependent and time-independent mistranslation covariates are equal (i.e. risk increase and risk decrease balance out for a net-zero risk) and predict that mistranslation, as a whole, begins to reduce risk of death at day 32 for V→S flies and day 25 for T→S flies. This predicted inflection timepoint occurs relatively early (before the 50% survival time) and flies that live beyond this timepoint have reduced risk when mistranslating compared to controls. We found that the interaction of mistranslation (time-independent covariate) and mating significantly decreased risk in female flies mistranslating from V→S by 1.81-fold compared to what would be expected based on the effects of mating and mistranslation individually (HR=0.551, *P*=0.0412; Supplemental Table S4), and a significant 2.24-fold decrease in risk for females mistranslating T→S (HR=0.445, *P*<0.0023; Supplemental Table S5), This indicates that the risk of death from mating and mistranslating is lessened when the two are combined, supporting the notion that mistranslation acts as a protective buffer against the negative effects of mating on longevity, and particularly so in longer-lived flies.

Lastly, it is possible that the increased longevity of mistranslating mated females is caused by these females not storing male ejaculate or having a reduced investment in reproduction due to energetic allocation towards resolving mistranslation over investing in reproduction (Priest *et al*. 2008; reviewed in Kressler *et al*. 2010; reviewed in Liu and Chern 2021). If mistranslating females do not bear the lifespan cost of reproduction because they are not storing sperm or investing in egg production. To assess whether mistranslating females are engaging fully in reproduction, we scored egg laying and fertilization to determine if mistranslation affects reproductive output. All mistranslating females and their controls produced eggs (oocytes) (Supplemental Table S3). *Drosophila melanogaster* females store sperm in the seminal receptacle for usage up to four days post-mating and the spermathecae for usage between five days to three weeks following mating (Dhillon *et al*. 2020, reviewed in Neubaum and Wolfner 1999). To determine if females produced the same amount of fertilized eggs from both sperm storage organs, we used a Linear Mixed Effects (LME) model on egg laying pattern of V→S females, T→S females, and their respective controls once mated to a wildtype (*CS*) male (Supplementary Table S3). Notably, there was a significant interaction effect between genotype and sperm storage period for V→S mistranslating females and their controls (*P*=0.0374; Table 3), which was not seen for T→S females. This interaction effect is due to the non-significant (via Wilcoxon test) larger and smaller amount of eggs produced from the short-term and long-term sperm storage organs of controls, respectively (Figure 5, Table 4). When short-term and long-term egg laying is pooled, there is no significant difference between V→S mistranslating females and their controls, but T→S mistranslating females lay significantly more eggs than controls (Wilcoxon test, *P=*0.018; Figure 6, Table 4). This due to a significant decrease in short-term egg laying of T→S control females (LME: *P*= 0.0334, Table 3; Wilcoxon test: *P*=0.018, Table 4).

**Figure 5.**
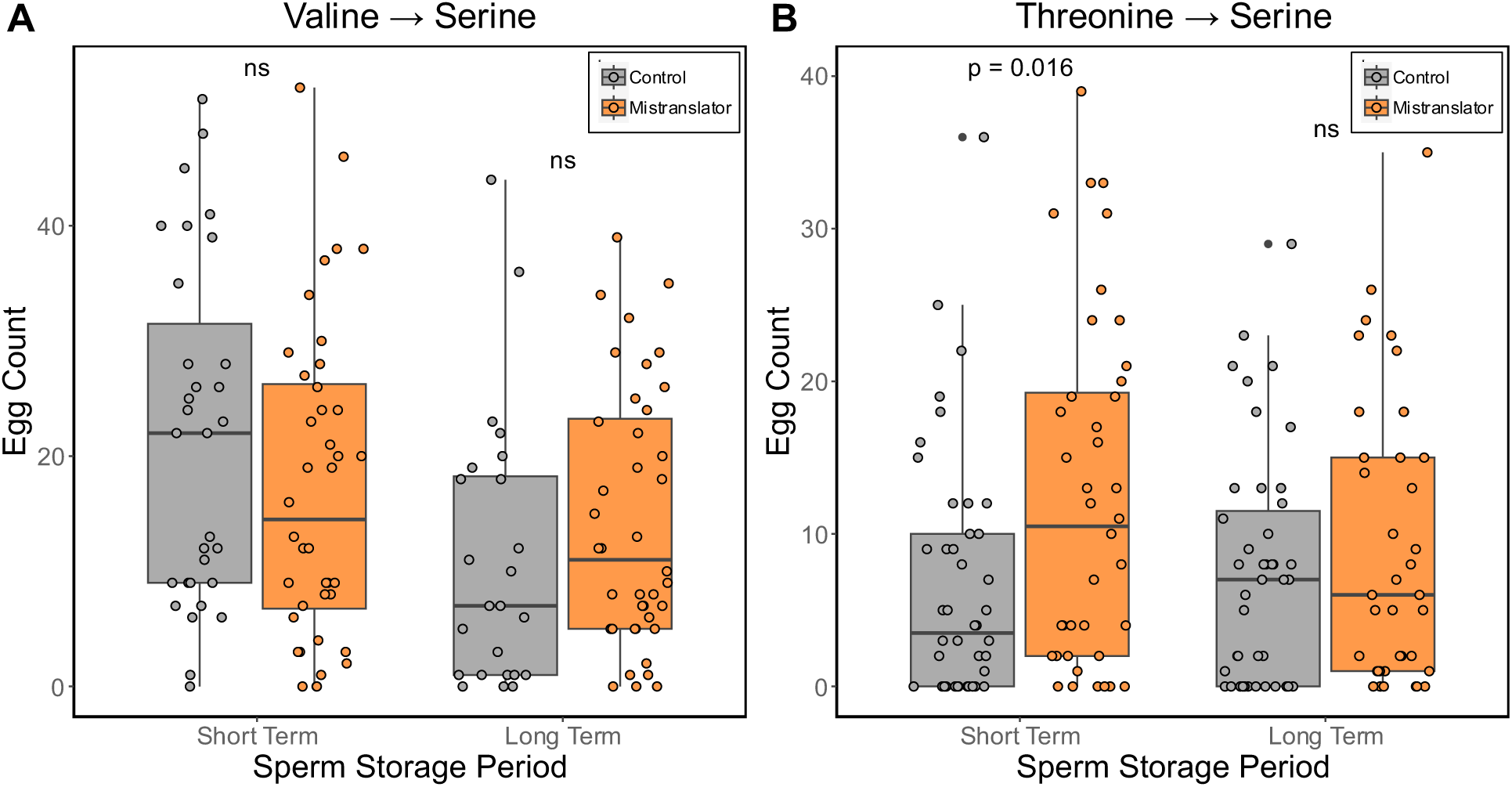
The number of eggs laid each day using sperm from short-term and long-term sperm storage organs for females with an a) V→S or b) T→S mistranslating tRNA (orange) compared to controls (grey) after mating with wildtype males. The boxes display the medians and interquartile ranges in the data distribution; circles represent individual samples.

**Figure 6.**
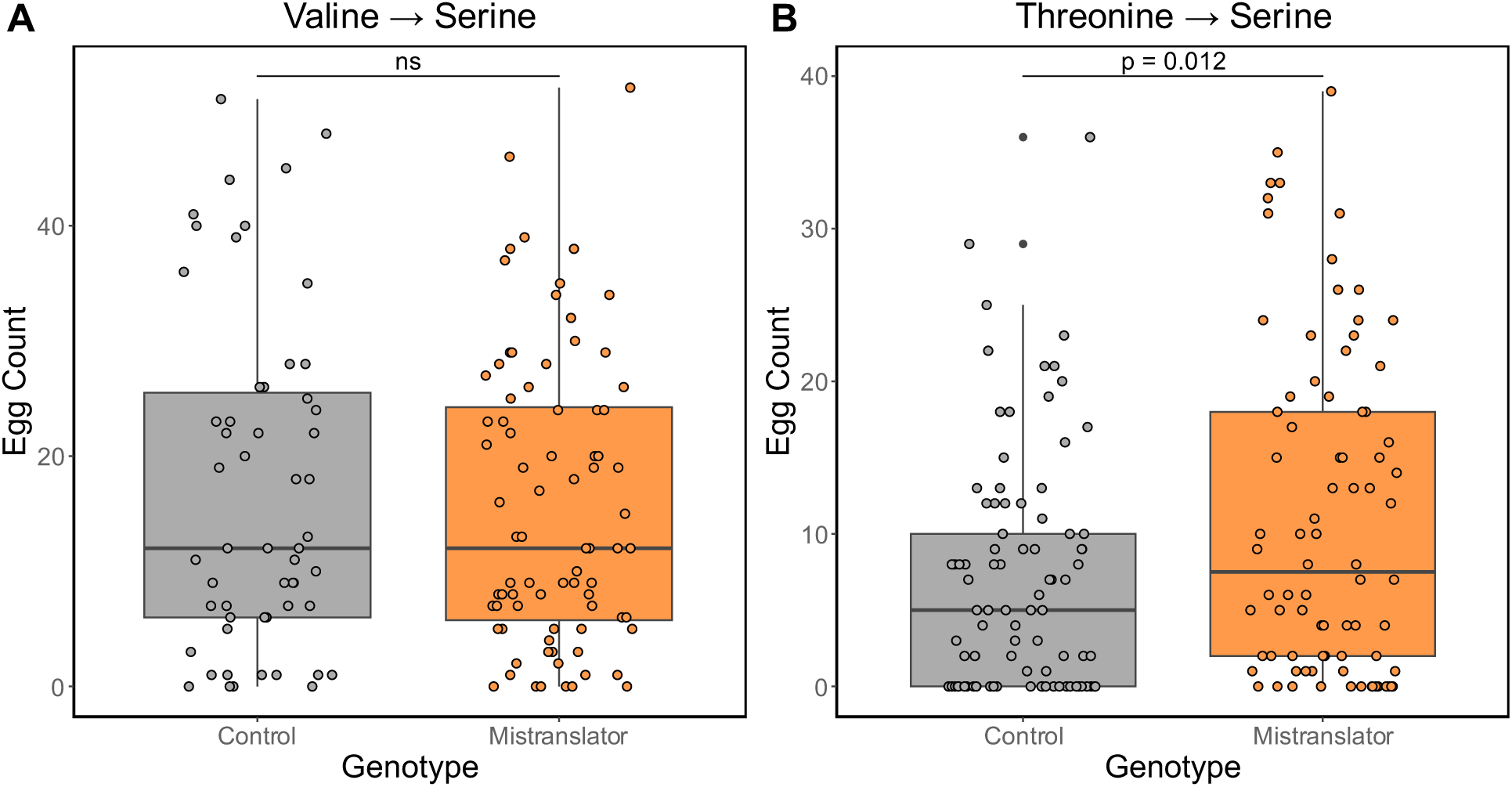
The number of eggs laid each day for females with an a) V→S or b) T→S mistranslating tRNA (orange) compared to controls (grey) after mating with wildtype males. The boxes display the medians and interquartile ranges in the data distribution; circles represent individual samples.

**Table 2.** Cox proportional hazard estimates and corrected *P-values* for all covariates in female V→S and T→S models.

| Model | Covariate<sup>1</sup> | Hazard ratio | *P*-value | Adjusted *P*-value<sup>2</sup> |
| --- | --- | --- | --- | --- |
| V→S females | mating status:mistranslation | 0.55060 | 0.01374 | **0.04122** |
|  | mating status | 4.06061 | 1.02E-14 | **3.05E-14** |
|  | mistranslation (time independent) | 4.61318 | 0.00348 | **0.00348** |
|  | mistranslation (time dependent) | 0.56557 | 7.40E-06 | **1.11E-05** |
|  | initial vial population size | 1.00938 | 0.78688 | 0.78688 |
| T→S females | mating status:mistranslation | 0.44527 | 0.00076 | **0.00227** |
|  | mating status | 4.80527 | 1.22E-18 | **3.66E-18** |
|  | mistranslation (time independent) | 10.66775 | 9.00E-06 | **2.70E-05** |
|  | mistranslation (time dependent) | 0.55873 | 3.81E-06 | **5.72E-06** |
|  | initial vial population size | 1.16577 | 0.00889 | **0.00889** |

<sup>1</sup> Reference levels: unmated (mating status), non-mistranslating control (mistranslation).
<sup>2</sup> *P*-value corrected for multiple testing using Benjamini-Hochberg false discovery rate. Significant values are bolded

**Table 3.** Linear mixed effects of egg laying and corrected *P-*values in V→S and T→S mistranslating and control females that were mated to wildtype males.

| Model | Covariate<sup>1</sup> | Covariate<sup>2</sup> | *P*-value |
| --- | --- | --- | --- |
| V→S females | – | storage | 0.2309 |
|  | storage | – | 0.1857 |
|  | storage:mistranslation |  | **0.0374** |
| T→S females | – | storage | **0.0334** |
|  | storage | – | 0.4147 |
|  | storage:mistranslation |  | 0.1711 |

<sup>1</sup>Reference levels: long term sperm storage (storage), non-mistranslating control (mistranslation)
<sup>2</sup>Reference levels: short term sperm storage (storage), non-mistranslating control (mistranslation)

**Table 4.** Wilcoxon test of egg laying in the short-term assay, long-term assay, and short-term and long-term assays pooled for V→S and T→S females that were mated to wildtype males.

| Model | Group 1 | Group 2 | Assay | *P*-value |  |
| --- | --- | --- | --- | --- | --- |
|  |  |  |  | Raw | Adjusted<sup>1</sup> |
| V→S females | Mistranslating | Control | Short | 0.17263 | 0.25895 |
|  |  |  | Long | 0.16684 | 0.25895 |
|  |  |  | Pooled | 0.77594 | 0.77594 |
| T→S females | Mistranslating | Control | Short | **0.00778** | **0.01752** |
|  |  |  | Long | 0.41792 | 0.41792 |
|  |  |  | Pooled | **0.01168** | **0.01752** |

<sup>1</sup>*P*-value corrected for multiple testing using Benjamini-Hochberg false discovery rate. Significant values are bolded

To confirm that eggs were fertilized, we tested for fertilization status halfway through short-term sperm storage (day 2 post-mating) and long-term sperm storage (day 8 post-mating) assays and found that 100% of eggs that were produced were fertilized. Results indicate that the increased longevity observed for mistranslating females is not due to a failure of sperm storage or lack of investment in reproduction, at least via the measures scored here.

## Discussion

According to the Disposable Soma Theory, reproduction leads to a reduction in lifespan due to the physiological cost of gamete production and favouring of gametic tissue proteostasis over somatic tissues (Kirkwood 1977; Evans *et al*. 1997; Kirkwood and Holliday 1997; reviewed in Flatt 2011). The cost of reproduction would be more pronounced in females than in males due to the higher energetic cost of female egg production than male sperm production. Based on the predictions of the Disposable Soma Theory, we predicted that a disruption to proteostasis due to mistranslation would further compound the detrimental effects of mating on lifespan. When we assessed the impact of reproduction and mistranslation on *Drosophila melanogaster* lifespan, we found that, counter to expectation, mistranslation caused by two different tRNA^Ser^ variants both increased mated female lifespan. There were no differences in egg laying between V→S mistranslating females and their controls, and T→S mistranslating females actually lay more eggs than their respective controls, further indicating that the increase in longevity observed cannot be attributed to a general failure or reduction in reproduction due to mistranslation. However, it is possible that other measures of reproductive investment, such as egg size, may reveal differences between mistranslating females and controls and warrant further examination. Mistranslation increased mated female lifespan to be equal to that of mated males, thus mistranslation appears to protect females from the detrimental impact of reproduction. Notably, mistranslation causes a synergistic and significant decrease in risk of death of female flies that are mistranslating and mated. Mistranslation therefore benefits female lifespan even when compounded with other stressors, indicating its potential as an evolutionary mechanism in multicellular organisms to improve adult fitness (Bacher *et al*. 2007; Netzer *et al*. 2009; Vargas-Rodriguez *et al*. 2021).

One limiting factor of our study is that we did not control for mating frequency during the longevity assay, which could impact longevity if different genotypes have different mating frequencies. For example, a higher mating frequency in early life has been shown to increase mortality in *D. melanogaster* females (Barnes *et al*. 2008; Travers *et al*. 2015). While male mortality was not different between the genotypes, and thus the number of males available to mate were similar over time, future studies that control for how often mating occurs throughout lifespan could determine if this is a contributing factor towards the longevity differences we observed. We also note that although we see significant effects due to mistranslation, we did not find the same increased longevity for T→S mistranslating females previously reported (Isaacson *et al*. 2024b) for these flies. It is possible that background genetic effects influencing longevity have arisen in this strain in the time between that earlier study and the one reported here, that the smaller sample size in the current study was insufficient to detect the T→S longevity increase, or that the effect on longevity is sensitive to subtle environmental differences that could be present between earlier assays and the ones here. Since T→S mistranslated at approximately twice the amount as V→S (1.0% *vs* 0.5% per codon, respectively; Isaacson *et al*. 2024b), then suppressors of mistranslation are more likely to have arisen in the T→S strain (Hoffman *et al*. 2017). If mistranslation is being partially suppressed in T→S flies to where it is removing the increased lifespan effect in unmated females, that would mean that our findings are likely under-representing the potential improved lifespan effects for mated females.

The removal of the impact of reproduction on lifespan in the presence of mistranslation is surprising. Why would mated female lifespan be longer with mistranslation than without? One mechanism could be heightened stress response pathways in the presence of mistranslation, causing the females to be ‘primed’ to respond to cellular stressors that arise with reproduction. A previous evaluation of gene expression with mistranslation in *Drosophila* did not find upregulation of stress-response genes (Isaacson *et al*. 2024), but stress response pathways are often increased at the protein level rather than transcript level (e.g., Kaarniranta *et al*. 1998; Harding et al. 2000; reviewed in Advani and Ivanov 2019). Stress response pathway levels may also be impacted by fragmentation of tRNAs, which can occur in response to stress, but whose effect would not be detected via sequencing of mRNAs. Fragmentation due to stressors depletes the amount of a tRNA available for translation (Huh *et al*. 2021), altering the tRNA pool in a way that causes increased levels of required proteins for different stress responses (Torrent *et al*. 2018). Thus, proteomic analysis of may identify some of the stress response pathways that differ between mated and unmated mistranslating flies.

Flies also have low levels of naturally occurring mistranslation (Isaacson *et al*. 2024b). When we induce substantial levels of mistranslation, this creates a strong selective pressure on the flies. Mistranslation from V→S and T→S leads to high rates of developmental mortality and physical abnormalities (Isaacson *et al*. 2024b). Flies with either the ability to suppress mistranslating tRNAs or to increase proofreading of protein production would have a higher chance of surviving to successfully reproduce. It may be that adults with a reduction of overall mistranslation, which could affect both transgenically-induced and naturally-occurring mistranslating tRNA variants, leads to increased longevity with particularly strong benefits during the protein-intensive process of reproduction. From a selection standpoint, positive selection for increased longevity due to the presence of a mistranslating tRNA would conflict with the negative effects of mistranslation during development. Additionally, there may be tissue-specific effects of mistranslation on lifespan that are not captured by the ubiquitous expression model used here. Therefore, further studies with mistranslation occurring only after completion of development, and assessment of the impacts of mistranslation confined to particular tissues, will help elucidate the underlying basis of the positive effect of mistranslation on lifespan.

## Data Availability Statement

Fly lines and plasmids are available upon request. The authors affirm that all data necessary for confirming the conclusions of the article are present within the article, figures, and Supplemental material. Supplementary Figure S1 and Tables S1-S5 are appended to the manuscript. Supplementary Table S6 contains all raw data. Supplementary File S7 contains the R code used to analyse all raw data.

## Supporting information

Supplemental Table S6 _ raw data

## Acknowledgements

We would like to thank Dr. Ben Ruben for his help with statistical analysis of the egg laying data and multiple *P-value* adjustment. We would also like to thank Dr. Joseane Moreira do Nascimento and Dr. Robert Cumming for helpful feedback on the project and Dr. Matthew Berg and Dr. Christopher Brandl for helpful comments on the manuscript.

---

**Supplemental Figure S1.**
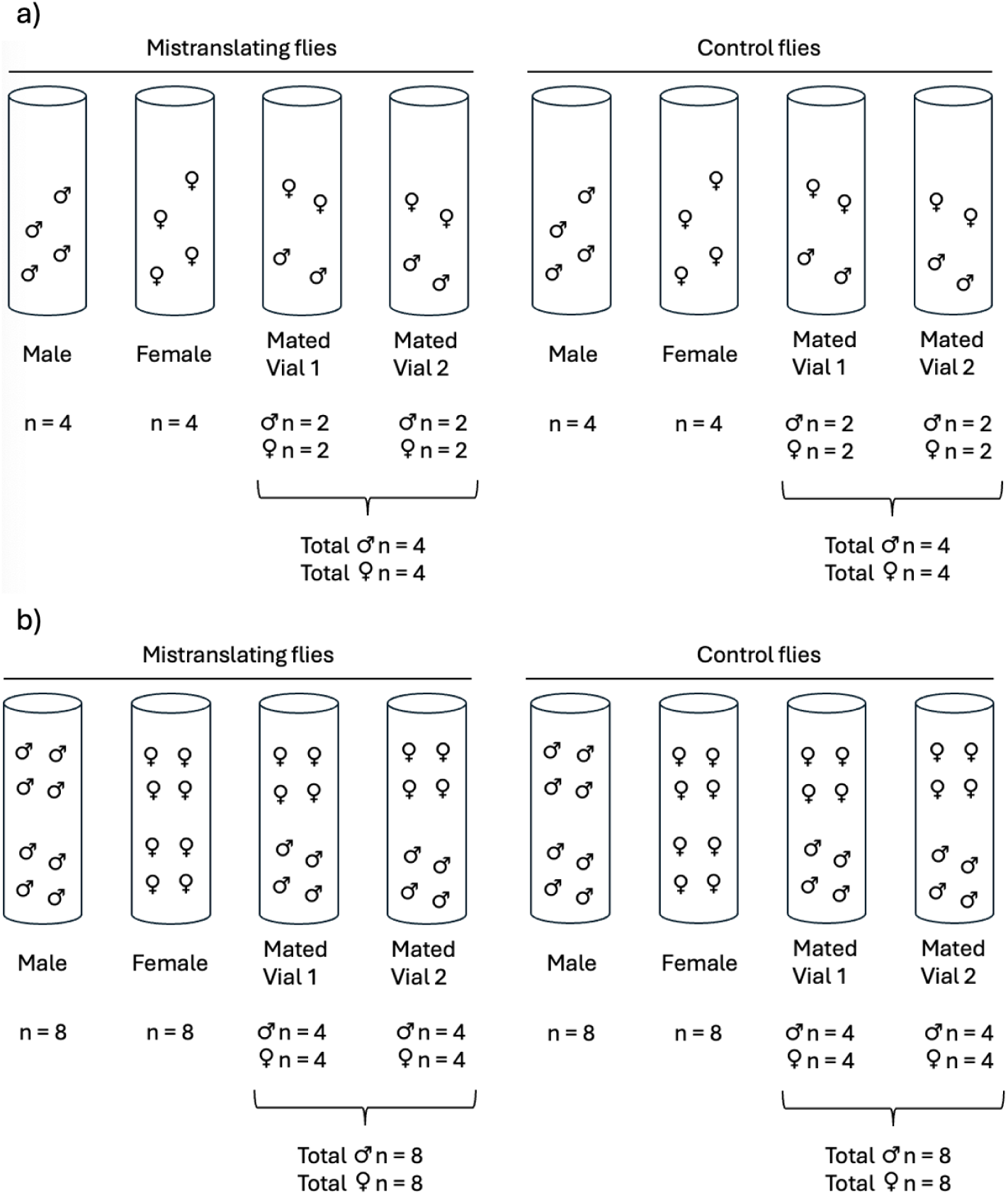
Longevity assay setup on a given day. Each male is depicted in the vials by the “♂” symbol and females are depicted by the “♀” symbol. The number of each fly in each vial is displayed beneath the vial. For a given set, vials contained a total of 4, 6, or 8 flies, with all vials within that set having the same number. The example displays a total of a) 4 flies and b) 8 flies.

**Supplemental Table S1.** Number of vials and flies in the longevity assay. The total number (n) of flies before censoring is shown, along with the number of data points that were censored.

| Model | Sex | Mating Status | Genotype | Vials | Flies | Censored Flies |
| --- | --- | --- | --- | --- | --- | --- |
| V→S | F | Unmated | Mistranlator | 18 | 80 | 2 |
|  |  |  | Control | 18 | 82 | 0 |
|  |  | Mated | Mistranlator | 36 | 82 | 0 |
|  |  |  | Control | 36 | 82 | 0 |
|  | M | Unmated | Mistranlator | 18 | 81 | 1 |
|  |  |  | Control | 18 | 81 | 1 |
|  |  | Mated | Mistranlator | 36 | 82 | 0 |
|  |  |  | Control | 36 | 81 | 1 |
| T→S | F | Unmated | Mistranlator | 16 | 76 | 4 |
|  |  |  | Control | 16 | 79 | 1 |
|  |  | Mated | Mistranlator | 32 | 79 | 1 |
|  |  |  | Control | 32 | 80 | 0 |
|  | M | Unmated | Mistranlator | 16 | 80 | 0 |
|  |  |  | Control | 16 | 80 | 0 |
|  |  | Mated | Mistranlator | 32 | 80 | 0 |
|  |  |  | Control | 32 | 80 | 0 |

**Supplemental Table S2.**
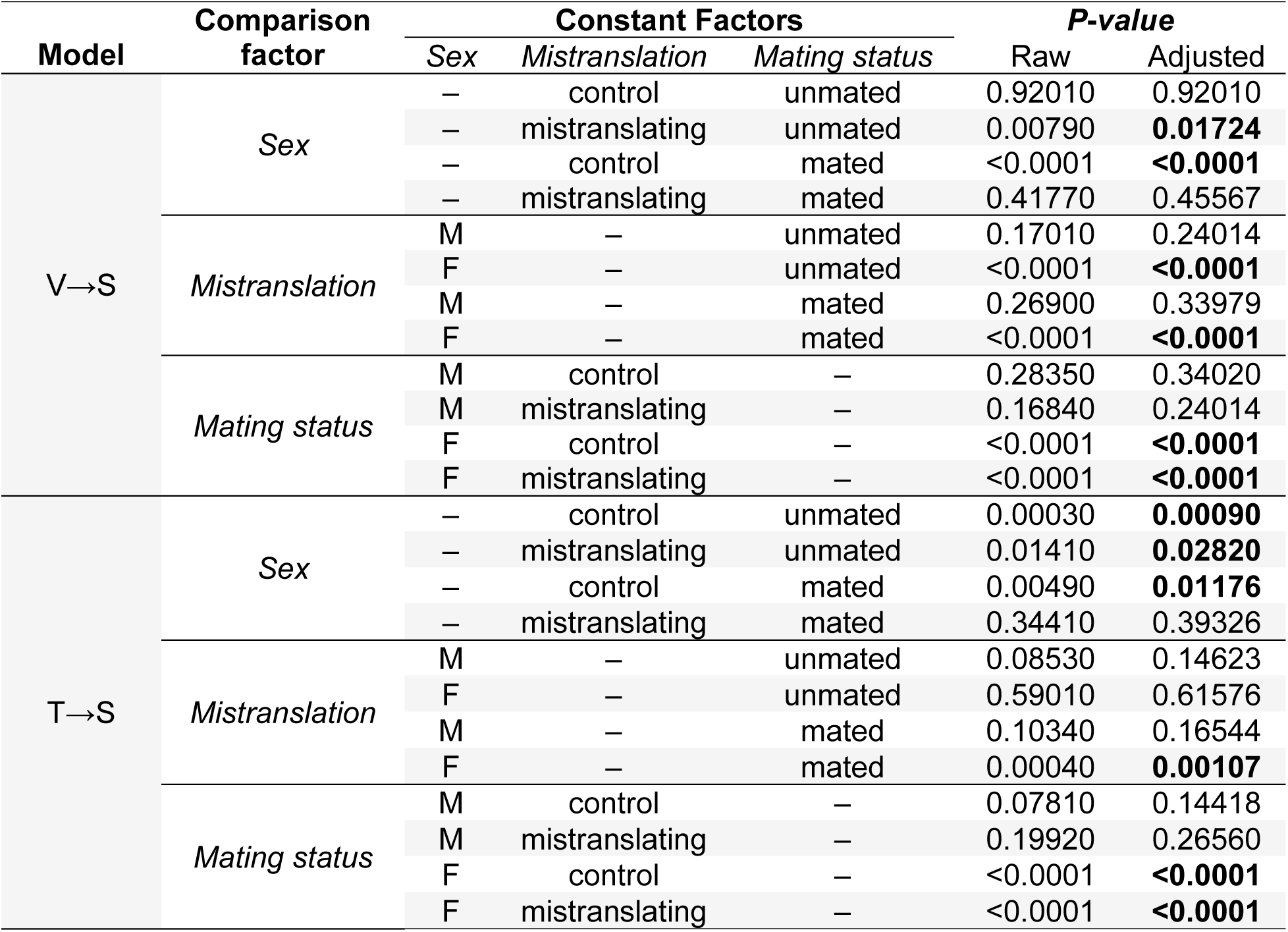
*P-*values of comparison groups. The table displays the unadjusted and BH-adjusted *P*-values for each mistranslating line (V→S and T→S) and their controls, categorised by sex (Males and Females) and mating status (mated and unmated). Each reference’s *P*-value is separated into their comparisons: Male x Female, MT x Control, and Mated x Unmated. Adjusted *P*-values that are bolded remained significant after adjustment.

| Model | Comparison factor | Constant Factors |  |  | *P*-value |  |
| --- | --- | --- | --- | --- | --- | --- |
|  |  | Sex | Mistranslation | Mating status | Raw | Adjusted |
| V→S | Sex | – | control | unmated | 0.92010 | 0.92010 |
|  |  | – | mistranslating | unmated | 0.00790 | **0.01724** |
|  |  | – | control | mated | <0.0001 | **<0.0001** |
|  |  | – | mistranslating | mated | 0.41770 | 0.45567 |
|  | Mistranslation | M | – | unmated | 0.17010 | 0.24014 |
|  |  | F | – | unmated | <0.0001 | **<0.0001** |
|  |  | M | – | mated | 0.26900 | 0.33979 |
|  |  | F | – | mated | <0.0001 | **<0.0001** |
|  | Mating status | M | control | – | 0.28350 | 0.34020 |
|  |  | M | mistranslating | – | 0.16840 | 0.24014 |
|  |  | F | control | – | <0.0001 | **<0.0001** |
|  |  | F | mistranslating | – | <0.0001 | **<0.0001** |
| T→S | Sex | – | control | unmated | 0.00030 | **0.00090** |
|  |  | – | mistranslating | unmated | 0.01410 | **0.02820** |
|  |  | – | control | mated | 0.00490 | **0.01176** |
|  |  | – | mistranslating | mated | 0.34410 | 0.39326 |
|  | Mistranslation | M | – | unmated | 0.08530 | 0.14623 |
|  |  | F | – | unmated | 0.59010 | 0.61576 |
|  |  | M | – | mated | 0.10340 | 0.16544 |
|  |  | F | – | mated | 0.00040 | **0.00107** |
|  | Mating status | M | control | – | 0.07810 | 0.14418 |
|  |  | M | mistranslating | – | 0.19920 | 0.26560 |
|  |  | F | control | – | <0.0001 | **<0.0001** |
|  |  | F | mistranslating | – | <0.0001 | **<0.0001** |

**Supplemental Table S3.** Number of eggs laid by each female for each cross after a single mating to a wildtype male. Days 1-4 represent short-term egg laying and days 7-10 represent long-term egg laying. NA represents flies that were harmed or lost during transfer. Note that T→S females #3 and #11 were removed from the analysis.

| Female | # | Number of Eggs Laid |  |  |  |  |  |  |  |
| --- | --- | --- | --- | --- | --- | --- | --- | --- | --- |
|  |  | Day 1 | Day 2 | Day 3 | Day 4 | Day 7 | Day 8 | Day 9 | Day 10 |
| V→S | 1 | 24 | 4 | 12 | 3 | 12 | 5 | 9 | 6 |
|  | 2 | 27 | 20 | 16 | 34 | 0 | 29 | 17 | 22 |
|  | 3 | 30 | 8 | 9 | 9 | 7 | 10 | 18 | 5 |
|  | 4 | 20 | 46 | 9 | 3 | 29 | 13 | 39 | 8 |
|  | 5 | 0 | 8 | 6 | 7 | 2 | 15 | 1 | 7 |
|  | 6 | 1 | 9 | 28 | 26 | 0 | 12 | 35 | 0 |
|  | 7 | 0 | 29 | 38 | 13 | 1 | 7 | 32 | 28 |
|  | 8 | 0 | 3 | 37 | 19 | 5 | 8 | 25 | 26 |
|  | 9 | 52 | 12 | 21 | 19 | 19 | 24 | 23 | 8 |
|  | 10 | 24 | 2 | 38 | 23 | 5 | 20 | 34 | 5 |
| V→S<br>Control | 1 | 40 | 13 | 23 | 28 | 23 | 10 | 19 | 5 |
|  | 2 | 45 | 7 | 7 | 9 | NA | NA | NA | NA |
|  | 3 | 39 | 0 | 1 | NA | NA | NA | NA | NA |
|  | 4 | 51 | 28 | 12 | 40 | 1 | 18 | 7 | 18 |
|  | 5 | 35 | 12 | 9 | 9 | 3 | 1 | 44 | 12 |
|  | 6 | 41 | 22 | 6 | 26 | 1 | 36 | 11 | 6 |
|  | 7 | 22 | 25 | 6 | 9 | 0 | 0 | 0 | 7 |
|  | 8 | 48 | 11 | 24 | 26 | 1 | 20 | 22 | 1 |
| T→S | 1 | 1 | 26 | 10 | 24 | 0 | 15 | 18 | NA |
|  | 2 | 31 | 20 | 4 | 13 | 0 | 2 | 6 | 2 |
|  | 3 | 6 | 32 | 10 | NA | NA | NA | NA | NA |
|  | 4 | 0 | 0 | 0 | 0 | 1 | 0 | 6 | 13 |
|  | 5 | 0 | 0 | 0 | 2 | 0 | 0 | 7 | 5 |
|  | 6 | 19 | 33 | 8 | 2 | 2 | 35 | 15 | 1 |
|  | 7 | 18 | 16 | 4 | 2 | 5 | 18 | 15 | 8 |
|  | 8 | 12 | 39 | 17 | 7 | 0 | 18 | 23 | 5 |
|  | 9 | 2 | 33 | 19 | 4 | 1 | 23 | 24 | 1 |
|  | 10 | 24 | 21 | 15 | 4 | 1 | 26 | 22 | 2 |
|  | 11 | 11 | 31 | 13 | 4 | 0 | 14 | 9 | 10 |
| T→S<br>Control | 1 | 0 | 12 | 5 | 10 | 0 | 0 | 0 | 0 |
|  | 2 | 19 | 9 | 5 | 2 | 7 | 13 | 8 | 8 |
|  | 3 | 1 | 9 | 0 | 0 | 0 | 9 | 0 | 21 |
|  | 4 | 4 | 8 | 3 | 0 | 0 | 2 | 17 | 8 |
|  | 5 | 4 | 0 | 0 | 5 | 2 | 7 | 13 | 0 |
|  | 6 | 16 | 12 | 9 | 7 | 0 | 0 | 6 | 12 |
|  | 7 | 2 | 0 | 0 | 0 | 0 | 0 | 7 | 8 |
|  | 8 | 0 | 0 | 0 | 0 | 0 | 2 | 5 | NA |
|  | 9 | 12 | 25 | 3 | 3 | 1 | 29 | 8 | 13 |
|  | 10 | 0 | 22 | 2 | 18 | 2 | 21 | 20 | 10 |
|  | 11 | 5 | 9 | 0 | NA | NA | NA | NA | NA |
|  | 12 | 0 | 15 | 36 | 10 | 11 | 23 | 8 | 18 |

**Supplemental Table S4.** Cox proportional hazard estimates and *P*-values for all covariates across sex-pooled, male, and female V→S survival models.

| V→S | Covariate<sup>1</sup> | Hazard ratio | *P*-value | Adjusted *P*-value<sup>2</sup> |
| --- | --- | --- | --- | --- |
| Interaction terms | mating status:sex:mistranslation | 0.43436 | **0.00901** | – |
|  | mating status:sex | 3.81052 | **5.01E-9** | – |
|  | sex:mistranslation | 0.66801 | **0.07002** | – |
|  | mating status:mistranslation | 1.19962 | 0.41414 | 0.41414 |
|  | – *male* | 1.20095 | 0.41366 | 0.41414 |
|  | – *female* | 0.55060 | **0.01374** | **0.04122** |
| Main terms | sex | 1.07818 | 0.63047 | – |
|  | mating status | 1.16402 | 0.33393 | 0.34872 |
|  | – *male* | 1.15945 | 0.34872 | 0.34872 |
|  | – *female* | 4.06061 | **1.02E-14** | **3.05E-14** |
|  | mistranslation (time independent) | 8.45394 | **9.04E-07** | **2.71E-06** |
|  | – *male* | 23.16539 | **0.00024** | **0.00037** |
|  | – *female* | 4.61318 | **0.00348** | **0.00348** |
|  | mistranslation (time dependent) | 0.54197 | **2.63E-09** | **7.88E-09** |
|  | – *male* | 0.42612 | **4.38E-05** | **0.00004** |
|  | – *female* | 0.56557 | **7.40E-06** | **0.00001** |
|  | initial vial population size | 1.05226 | **0.03882** | 0.05823 |
|  | – *male* | 1.10624 | **0.00448** | **0.01345** |
|  | – *female* | 1.00938 | 0.78688 | 0.78688 |

<sup>1</sup>Reference levels: unmated (mating status), male (sex), non-mistranslating control (mistranslation).
<sup>2</sup>*P*-values corrected for multiple testing for all one-way and two-way interactions using Benjamini-Hochberg false discovery rate. Significant values are bolded.

**Supplemental Table S5.** Cox proportional hazard estimates and *P*-values for all covariates across sex-pooled, male, and female T→S survival models.

| T→S | Covariate<sup>1</sup> | Hazard ratio | *P*-value | Adjusted *P*-value<sup>2</sup> |
| --- | --- | --- | --- | --- |
| Interaction terms | mating status:sex:mistranslation | 0.58387 | 0.09459 | — |
|  | mating status:sex | 3.09235 | **8.10E-07** | — |
|  | sex:mistranslation | 1.47627 | 0.08673 | — |
|  | mating status:mistranslation | 0.82055 | 0.37985 | 0.47595 |
|  | — *male* | 0.85158 | 0.47595 | 0.47595 |
|  | — *female* | 0.44527 | **0.00076** | **0.00227** |
| Main terms | sex | 0.51179 | **0.00003** | — |
|  | mating status | 1.41349 | **0.02888** | **0.03343** |
|  | — *male* | 1.40130 | **0.03343** | **0.03343** |
|  | — *female* | 4.80527 | **1.22E-18** | **3.66E-18** |
|  | mistranslation (time independent) | 4.51845 | **0.00003** | **0.00005** |
|  | — *male* | 2.74026 | **0.03315** | **0.03315** |
|  | — *female* | 10.66775 | **0.00001** | **0.00003** |
|  | mistranslation (time dependent) | 0.62727 | **6.56E-08** | **1.97E-07** |
|  | — *male* | 0.72124 | **0.00625** | 0.00625 |
|  | — *female* | 0.55873 | **3.81E-06** | **0.00001** |
|  | initial vial population size | 1.19454 | **0.00002** | **0.00005** |
|  | — *male* | 1.21652 | **0.00071** | **0.00106** |
|  | — *female* | 1.16577 | **0.00889** | **0.00889** |

<sup>1</sup>Reference levels: unmated (mating status), male (sex), non-mistranslating control (mistranslation).
<sup>2</sup>*P*-values corrected for multiple testing for all one-way and two-way interactions using Benjamini-
Hochberg false discovery rate. Significant values are bolded.

